# Low entropy map of brain oscillatory activity identifies spatially localized events: a new method for automated epilepsy focus prediction

**DOI:** 10.1101/707497

**Authors:** Manel Vila-Vidal, Carmen Pérez Enríquez, Alessandro Principe, Rodrigo Rocamora, Gustavo Deco, Adrià Tauste Campo

## Abstract

The spatial mapping of localized events in brain activity critically depends on the correct identification of the pattern signatures associated with those events. For instance, in the context of epilepsy research, a number of different electrophysiological patterns have been associated with epileptogenic activity. Motivated by the need to define automated seizure focus detectors, we propose a novel data-driven algorithm for the spatial identification of localized events that is based on the following rationale: the distribution of emerging oscillations during confined events across all recording sites is highly non-uniform and can be mapped using a spatial entropy function. By applying this principle to EEG recording obtained from 67 distinct seizure epochs, our method successfully identified the seizure focus on a group of ten drug-resistant temporal lobe epilepsy patients (average sensitivity: 0.94, average specificity: 0.90) together with its characteristic electrophysiological pattern signature. Cross-validation of the method outputs with postresective information revealed the consistency of our findings in long follow-up seizure-free patients. Overall, our methodology provides a reliable computational procedure that might be used as in both experimental and clinical domains to identify the neural populations undergoing an emerging functional or pathological transition.

**Conflict of interests:** The authors declare no competing financial interests.

## Introduction

The study of high cognitive functions or brain diseases with electroencephalography often involves the identification of changes in the recorded potentials that are time-locked to an event (Luck et al., 2000; Lauchaux et al., 2012; Kropotov, 2016). Depending on the nature of the problem being addressed, this event can be the onset of an external stimulus, a motor act, or a pathological symptomatology. In all cases, the spatial localization of the changes elicited in brain activity depends on the identification of the pattern signatures of those changes.

For instance, in the context of epilepsy research and monitoring with intracranial EEG, clinicians target specific electrophysiological patterns that are known to be associated with epileptic activity. In particular, the spatial mapping of pathological patterns of activity in drug-resistant epilepsy patients undergoing pre-surgical stereo-electroencephalography (SEEG) (Talairach et al., 1974; Munari and Bancaud, 1985; Guenot et al., 2002; Engel et al., 2005) is a crucial step to delineate the seizure onset zone (SOZ) and plan a successful surgery.

Over the last decades, the problem of seizure focus localization from intracranial EEG recordings has fostered the development of quantitative tools to better characterize and understand ictal genesis and propagation. Several biomarkers characterize the epileptogenicity of the monitored brain structures based on preselected spectral features of the signal (Bartolomei *et al*., 2008; David *et al.*, 2011; Gnatovsky *et al*., 2011, 2014; Andrzejak *et al*., 2015; Vila-Vidal *et al*., 2017). More recent studies have proposed automatic methods based on high-frequency oscillations (HFOs) or stochastic properties of the SEEG signals in predefined frequency windows (Geertsema *et al*. 2015; Liu *et al*., 2016; Murphy *et al*., 2017; Varatharajah *et al*., 2017).

Despite many efforts, the gold standard in clinical practice still remains visual inspection of EEG recordings due a number of reasons. For instance, the heterogeneity of electrophysiological patterns associated with seizure onset (Perucca et al., 2014; Lagarde et al., 2016) represents a major drawback to design SOZ detection algorithms that are universally valid for all seizure typologies and patients. All cited works made a priori assumptions on the spectral features of ictal-driven activity (e.g. using HFOs or power in the broadband spectrum) and might turn ineffective for seizures that don’t fulfil such frequency constraints. For example, SOZ detectors based on HFOs are very specific to fast discharges, but might overlook pathological patterns of activity dominated by lower frequency oscillations and longer temporal scales.

In the present study, we propose a novel data-driven methodology for the spatial localization of spatially confined events in brain activity from intracranial EEG recordings that makes no assumption on the spectral properties of the pattern signatures of those events. Our method relies on finding the temporal scale and frequency range of locally enhanced neural oscillations associated with the event of interest. Central to our method is the definition of two novel measures, the global activation (GA) and the activation entropy (AE), that quantify the magnitude of spectral changes with respect to a pre-defined baseline and the spread of these activations across recording sites, respectively, at different frequencies and as time progresses from the occurrence of the event. By setting appropriate conditions on these measures, it is possible to find time-frequency windows where the most relevant sites can be optimally discriminated.

To validate our algorithm, we adjusted and applied it to peri-ictal SEEG recordings to identify the seizure onset patterns and localize the seizure onset zone (SOZ). In this context, spatially confined activations with respect to a pre-ictal baseline were taken as signatures of epileptogenic tissue and were used to localize the seizure focus. The algorithm’s performance on a group of ten patients with varied seizure onset patterns and follow-up periods that range from 3 to 6 years. In each case, the underlying time-frequency windows were found to pinpoint the characteristic spectral features (frequency and duration) of onset patterns. Additionally, we were able to relate our findings with the postsurgical information of patients that attained seizure freedom after resective surgery using the core measure of the study as a putative predictor of the resected zone. In the hope that it will be useful to other researchers, some of the processing tools used in this work have been publicly released as an open-access Python package (Epylib v1.0: https://github.com/mvilavidal/Epylib/wiki).

Overall, the proposed procedure could be used in a variety of settings to extract the pattern signatures and spatial localization of the brain response to events of interest in a variety of settings of both clinical and cognitive nature. For example, in finding the brain sites that respond to electrical stimulation or to drug administration, in the analysis of the brain reaction to cognitive stimuli during a task paradigm, in studying the increase in connectivity between distant regions when a stimulus is delivered, etc. In particular, our classifier is particularly suitable to be used as a complementary tool during the pre-surgical evaluation and planning to better identify and interpret the regions involved in seizure generation and propagation.

## Materials and methods

The method described here can be used to extract the pattern signatures associated to localized events of clinical or cognitive interest recorded with the use of electroencephalography techniques. Without loss of generality, our method was validated in the SOZ localization setting, where a well-established ground truth is available for comparison. For the sake of simplicity, the methods’ description will be referred to the case of seizure focus localization from peri-ictal SEEG recordings (Fig. 1A,B).

**Figure 1.**
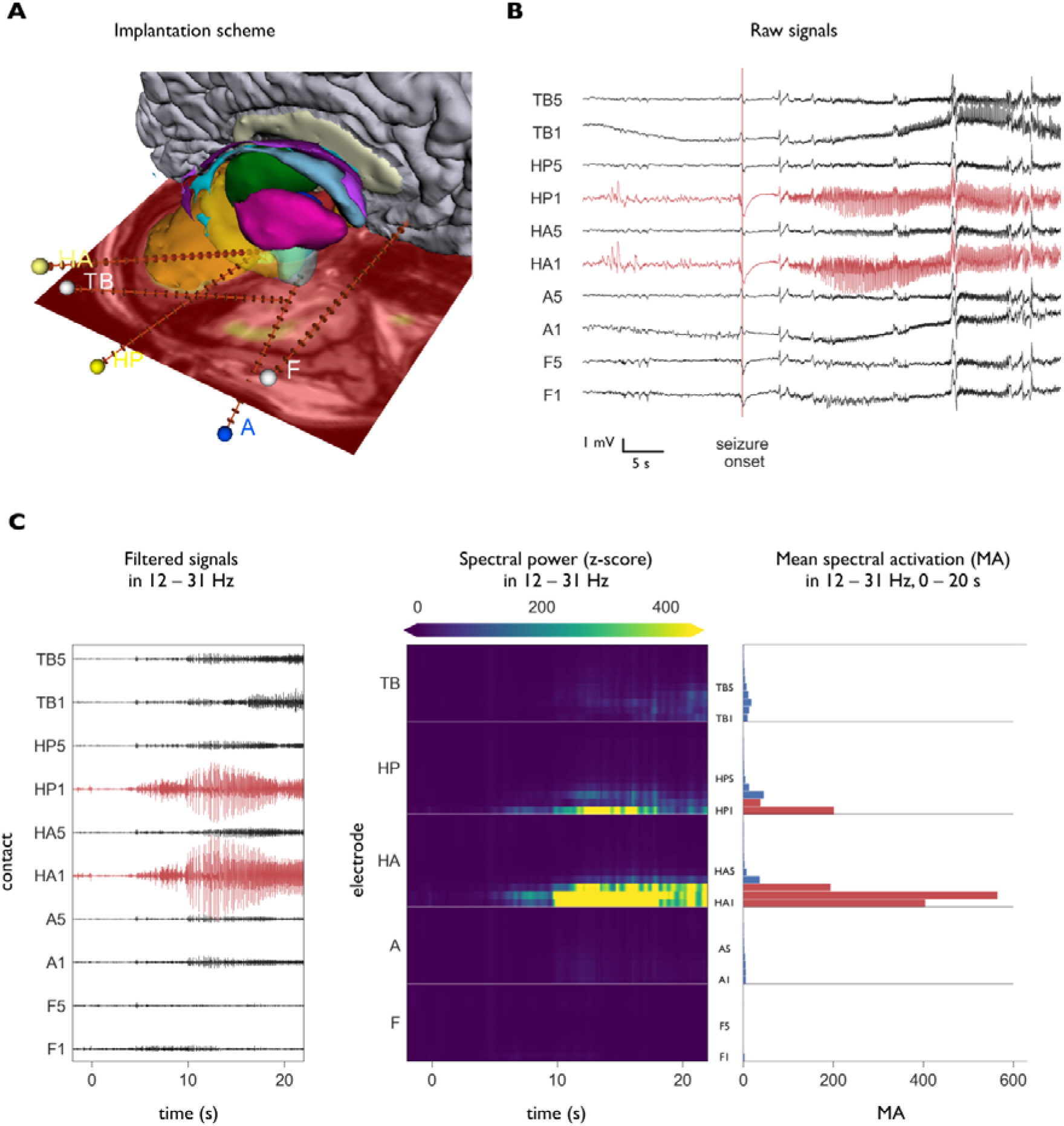
Example of SEEG signal analysis for a time window (0-20 seconds after seizure onset) and frequency band of interest (12-31 Hz) in a seizure of patient 1. (A) Medial view of the brain showing the trajectories of the five electrodes and their target regions. Electrode labels refer to the regions monitored by their internal contacts: lateral parts of the orbitofrontal cortex (F), amygdala (A), anterior hippocampus (HA), posterior hippocampus (HP) and temporal pole (TB). Contacts within an electrode are numbered starting from inside. Contacts HA1, HA2, HA3, HP1 and HP2 were identified as being part of the seizure onset zone in the pre-surgical evaluation. (B) SEEG recordings from a selection of contacts (two per electrode) around the seizure onset (20 seconds of pre-ictal and 40 seconds of the seizure period are shown). SOZ contacts are marked in red. (C) Signal analysis in a given time window (0-20s after SO) and frequency band (12-31 Hz) of interest. Signals were first band-pass filtered at 12-31 Hz and the first 20s of ictal activity were selected for analysis (left). Each contacts’ instantaneous spectral power was obtained using the Hilbert transform method and z-scored with respect to a baseline distribution defined by accumulating the power values of all contacts during the first 40 seconds of the pre-ictal period (middle). The mean activation (MA) was computed for each contact by averaging all power values in the first 20 s of the ictal period (right).

### Low entropy map of enhanced neural oscillations

#### Magnitude and spread of enhanced oscillations: Global activation (GA) and Activation entropy (AE)

The current approach builds upon the previously introduced mean activation (MA) measure (Vila-Vidal *et al*., 2017), which quantifies the average spectral activation of each targeted brain structure with respect to a certain baseline period for pre-defined frequency and time windows of interest (Fig. 1C). In our case, we use a pre-ictal baseline of activity (from 60 to 20 seconds before ictal onset). A detailed description of the computation of the MA can be found in the Supplementary Information. We will denote the MA of a given region *j* in the frequency band *f* and computed over a time window spanning from the seizure onset until time *t* with the following notation: *MA_j_*(*f,t*).

For optimal focus detection we must ensure that there is a hierarchical and selective activation of SOZ contacts only. Central to our approach is the definition of two novel measures, namely the global activation (GA) and the activation entropy (AE), that are jointly optimized to find time-frequency windows of interest where ictal activity is maximal with respect to a baseline pre-ictal period, and is spatially confined to a few contacts. Fig. 2 illustrates how these measures are computed and used to assess the amount of information carried in each window of interest. On one hand, the global activation (GA) quantifies the magnitude of the most relevant spectral activations with respect to the pre-ictal baseline state for a given time-frequency window of interest. It is defined as the weighted average MA over all contacts, where each contact’s contribution is weighted by its own activation, thus ensuring that most active regions have a higher impact on the final value (Fig. 2A):

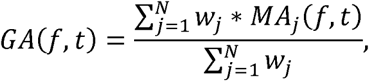

with *w_j =_ MA_j_ (f,t)* if *MA_j_ (f,t)*> 0, and *w_j_* = 0, if *MA_j_ (f,t)* ≤ 0. On the other hand, the AE is defined as the entropy of the MA distribution and characterizes the spatial spread of spectral activations for the given time-frequency of interest. First, the MA histogram is computed using 10 bins homogeneously spaced between the minimum and maximum MA values. Probability values for each bin (*p*_i_ for *i* = 1, …, 10) are found as the fraction of contacts lying within the corresponding MA bin (Fig. 2A). We then compute the Shannon’s entropy of this distribution using the formula:

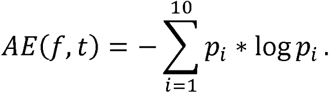

**Figure 2.**
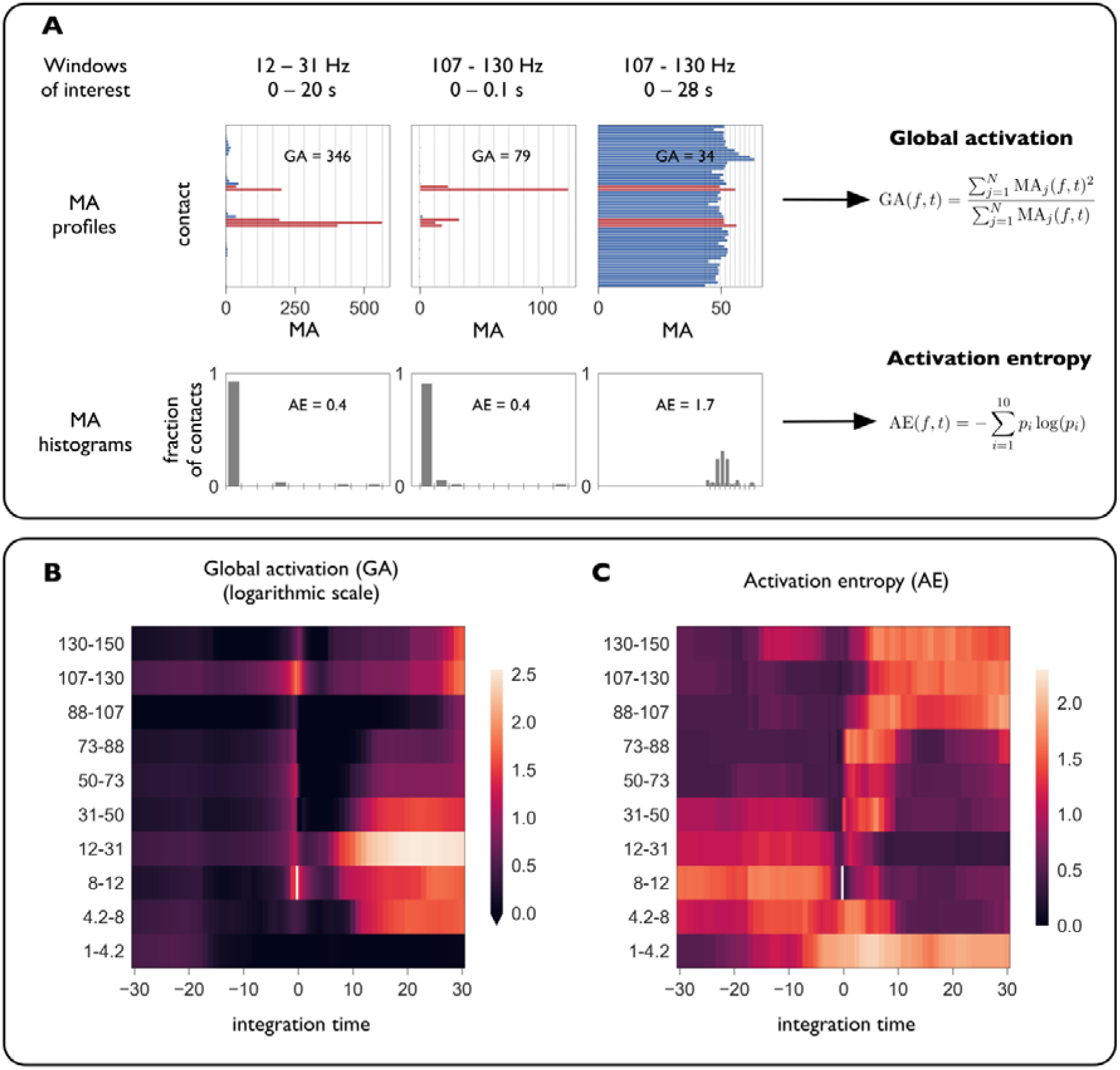
Two novel measures characterize the magnitude and spread of enhanced neural oscillations. Exploration of spectral activations in different frequency and time windows of interest is done around seizure onset. (A) MA profiles (mean spectral activation) and MA histograms (with 10 bins) for an exemplary selection of windows of interest. Visually marked SOZ channels are shown in red. The first two time-frequency windows (12-31 Hz, 0-20 s; 107-130 Hz, 0-100 ms) result in very structured and selective activations of SOZ channels. The third combination (107 - 130 Hz, 0-28 s) shows a uniform channel activation, indicating that averaged high-frequency activity is present in all recording sites. Two measures characterize the suitability of a given time-frequency window for SOZ detection. The global activation (GA) quantifies the magnitude of the highest activations for a given frequency and integration time. It is defined as the weighted average MA over contacts, where each contact’s contribution is weighted by its MA value (most active channels have a higher impact on the final value). The activation entropy (AE) quantifies how spread spectral activations are across recording sites. It is obtained by computing the Shannon’s entropy of the MA histogram with 10 bins homogeneously spaced between the minimum and maximum MA values. Lower entropies of the MA distribution indicate structured and spatially confined spectral activations, whereas higher values indicate distributed and spatially extended activations. (B) Global activation (GA) across all possible combinations of time and frequency windows of interest in the peri-ictal period. Positive and negative times denote time windows spanning from seizure onset into the ictal and pre-ictal epochs, respectively. (C) Activation entropy (AE) across all possible combinations of time and frequency windows of interest in the peri-ictal period.

In order to systematically explore different windows for optimal SOZ detection we first divided the frequency spectrum into 10 non-overlapping bands using the following cutting points: 1, 4.2, 8, 12, 31, 50, 73, 88, 107, 130 and 150 Hz. In each band we considered a set of 75 different integration time windows obtained by fixing the initial bound at the seizure onset and varying the final bound from 0.1 to 30 s (in steps of 0.1 s in the range 0.1-5 s and steps of 1 s in the range 5-30 s). Additionally, we also considered time windows spanning from seizure onset into the pre-ictal period, i.e., with final bound at seizure onset and initial bound ranging from 30 s to 0.1 s before seizure onset, with the same spacing as before. For each time-frequency window, we computed the MA profile and extracted its GA and AE. Fig. 2B and 2C show the GA and AE distributions for one exemplary seizure of patient 1, respectively.

#### Pattern signature and spatial map of localized emerging oscillations

The procedure described in the previous section was sequentially applied to the 67 seizures included in this study. Fig. 3 summarizes the processing steps. SEEG signals in the peri-ictal period were band-pass filtered in pre-defined bands of interest spanning the whole spectrum (Fig. 3A). Then, for each band, MAs were obtained for all possible time windows of interest (Fig. 3B). For each time-frequency window, we extracted the GA and AE from the MA distribution (Fig. 3C).

**Figure 3.**
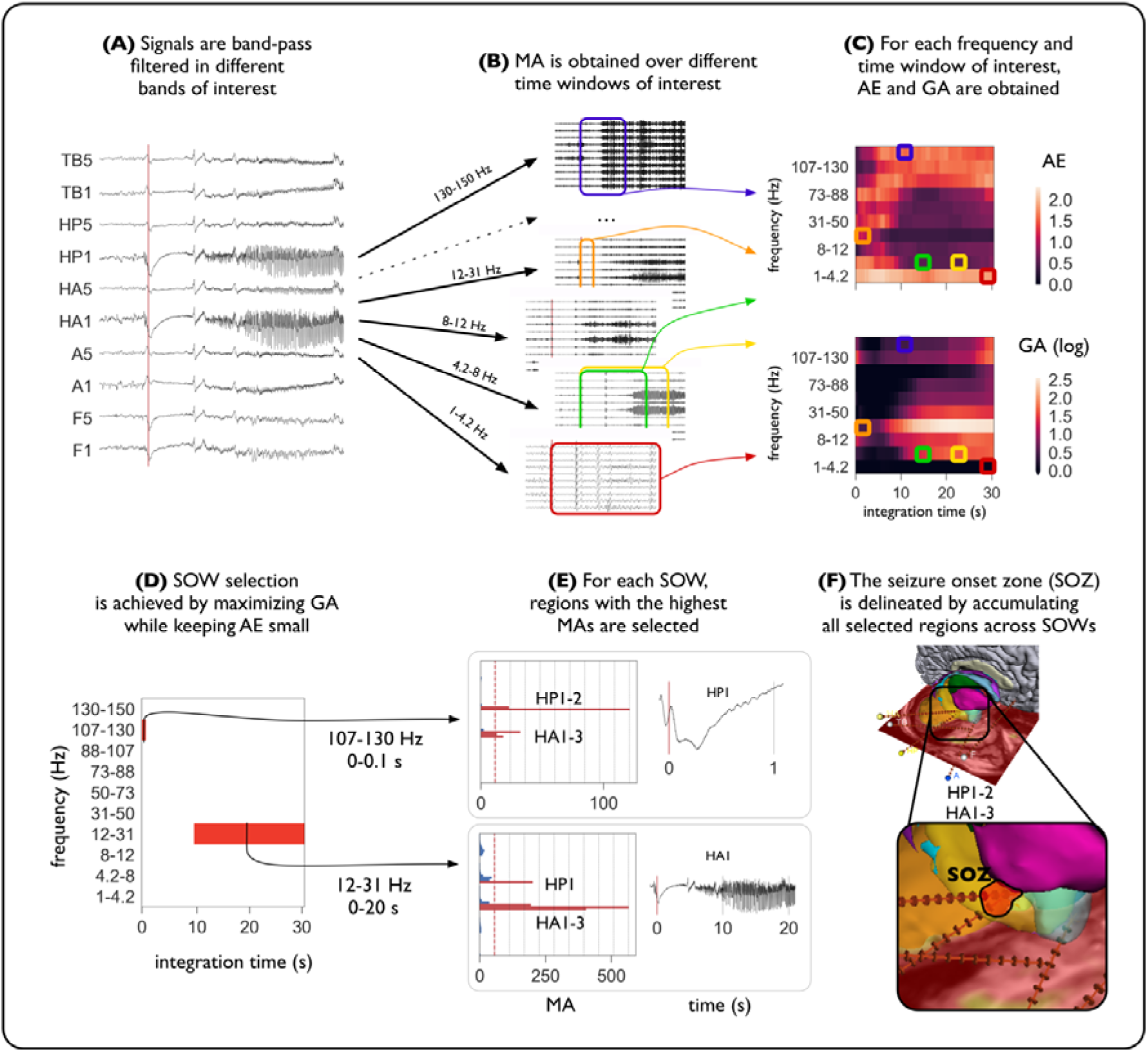
Processing steps from SEEG broadband signals to SOZ detection. Identification of the most relevant seizure onset windows (SOW) and seizure onset zone (SOZ) detection is illustrated with the first seizure of patient 1. (A) Signals are band-pass filtered in pre-defined bands of interest spanning the whole spectrum. (B) The mean spectral activation (MA) is obtained over different time windows and frequencies of interest using the Hilbert transform method and averaging instantaneous spectral activations across time. (C) For each time-frequency window, the MA profile is characterized by two summary measures: the global activation (GA) and the activation entropy (AE). (D) SOW detection is achieved by finding time-frequency windows that maximize GA under the constraint of low AE. Frequencies and time windows for which GA is above the 95th percentile and the AE is below 0.5 are selected. Selected time-frequency windows are marked in red. (E) For each seizure onset window, regions with MAs above the AE-induced threshold (dotted red line) are considered to be part of the SOZ. Active regions are marked in red in the bar plots. (F) Active regions are accumulated across all SOWs, thus obtaining a single SOZ for each seizure (shaded in red).

Seizure onset window (SOW) detection was achieved by finding time-frequency windows that maximized the GA under the constraint of low AE to ensure that spectral activations were confined only to a few contacts. We considered all pairs (f,t) with positive integration times (i.e., excluding time windows in the pre-ictal state) and set two threshold conditions, one per variable. For each seizure, we set a GA threshold at the 95-th percentile of the GA distribution. In any case, the threshold was always kept larger than 3 to ensure significant global activations with respect to the pre-ictal state. On the other hand, we set a threshold of 0.5 on the AE, which requires at least 80% of the contacts to lie within the same MA bin (see Supplementary Information). Specifically, they were required to lie within the lowest MA bin. All time-frequency windows satisfying both conditions were preselected as candidates to be SOWs. Finally, for each frequency band we kept only the first time-windows satisfying the condition. Fig. 3D shows the selected SOWs for the first seizure of patient 1.

As a result, at least 80% of the contacts are confined to the first bin of the MA histogram, exhibiting very low activations. The few remaining active contacts were considered to be part of the SOZ (Fig. 3E). This procedure was repeated for all selected SOWs, and SOZ contacts were accumulated, thus obtaining a single SOZ per seizure (Fig. 3F).

### Seizure focus prediction

#### Ethics statement

All diagnostic and surgical procedures have been previously approved by The Clinical Ethical Committee of Hospital del Mar (Barcelona, Spain).

#### Patient selection

We selected ten patients with pharmacoresistant temporal lobe epilepsy that underwent stereo-EEG in the Epilepsy Monitoring Unit of Hospital del Mar (Barcelona, Spain). Patient inclusion was based on the following criteria: a) that the seizure focus had been identified by the epileptologists and b) that ictal onset was confined to a reduced number of contacts corresponding to an anatomical region. Demographic and clinical information for all patients is summarized in Table 1.. The decision to implant depth electrodes, the decision of the targeted areas and the implantation duration were based solely on clinical grounds and independent of this study.

**Table 1.** Main data of patients included in the study. F = female; M = male; TLE=temporal lobe epilepsy; R=right; L=left; FC=frontal cingulate; R FC=right frontal cingulate; L FC=left frontal cingulate; A=amygdala; Ha=anterior hippocampus; Hp=posterior hippocampus; TP=temporal pole; EC=entorhinal cortex, OFCm=mesial orbitofrontal cortex; TGi=inferior temporal gyrus; PHCp=posterior parahippocampal cortex; W=Wernicke?s area; TOJ=temporal occipital junction; FBC=frontal basal cortex; MS=motor strip; TCl= lateral temporal cortex; OCm=mesial occipital cortex; FS=focal seizure; w=with; wo=without; CA=consciousness alteration; TATL: Tailored anterior temporal lobectomy; RF-TC=Radiofrequency thermocoagulation; SAH=Selective amygdalohyppocampectomy; Rp=Responsive; uRp=Unresponsive; NO=not-operated. *Patient 5 was initially responsive to RF-TC, but experienced seizure relapse two years after the procedure. He then underwent resective surgery, after which achieved seizure freedom (Engel I) with a follow-up of 3 years.

| Patient | Gender/<br>Age | Epilepsy | Side | Duration<br>(years) | Electrodes<br>(left) | Number of<br>recording<br>sites | Number<br>of sites<br>within<br>the SOZ | Analyzed<br>seizures | MRI | Surgery | Treated structure | Outcome<br>(Engel's<br>class) | Follow-up<br>(years) |
| --- | --- | --- | --- | --- | --- | --- | --- | --- | --- | --- | --- | --- | --- |
| 1 | F/27 | TLE | R | 10 | 5(0) | 54 | 5 | 8 | negative | R TATL | Temporal pole, amygdala and<br>head of hippocampus | Ia | 6 |
| 2 | F/30 | TLE | L | 25 | 7(7) | 67 | 9 | 8 | negative | SAH | temporal pole, inferior half of<br>amygdala and anterior 1/3 of<br>hippocampus | Ic | 6 |
| 3 | F/55 | TLE | L | 40 | 6(6) | 59 | 6 | 1 | negative | L TATL | temporal pole, amygdala and<br>head of hippocampus | Ib | 6 |
| 4 | M/29 | TLE | R | 9 | 0(0) | 95 | 11 | 8 | negative | NO |  | Ia | 5 |
| *5 | M/43 | TLE | R | 42 | 15(0) | 125 | 3 | 5 | FCDIIIa.<br>Arachnoid Cyst. | R TATL | temporal pole, amygdala and<br>2/3 of hippocampus | Ia | 3 |
| 6 | M/26 | TLE | R | 11 | 7(0) | 78 | 11 | 7 | R amygdala enlargement | RF-TC | temporal pole, amygdala, entorhinal cortex and<br>fusiform gyrus | Rp (Ib) | 4 |
| 7 | M/20 | TLE | L | 12 | 15(15) | 122 | 6 | 7 | negative | RF-TC | anterior and posterior<br>superior temporal gyrus,<br>transverse temporal gyrus<br>and supramarginal gyrus | Rp (Ib) | 4 |
| 8 | M/40 | TLE | L | 2 | 8(8) | 85 | 5 | 7 | L temporal polar<br>blurring | RF-TC | temporal pole and<br>hippocampal head | uRp (III) | 4 |
| 9 | F/52 | TLE | L | 45 | 10(8) | 104 | 8 | 6 | gliosis near R<br>craniotomy | L TATL | temporal pole, inferior half of<br>amygdala and anterior 1/3 of<br>hippocampus | Ib | 5 |
| 10 | F/40 | TLE | L | 16 | 10(8) | 107 | 14 | 10 | L posterior<br>hippocampal<br>lesion | L TATL | 2/3 of amygdala, anterior 2/3<br>of hippocampus | III | 6 |

#### SEEG recordings

Recordings were obtained using 5 to 21 intracerebral multiple contact microelectrodes (Dixi Medical, Besançon, France; diameter: 0.8 mm; 5 to 15 contacts, 2 mm long, 1.5 mm apart) that were stereotactically implanted using robotic guidance (ROSA, Medtech Surgical, Inc). Between 56 and 126 contacts were implanted and recorded in each subject. Fig. 1A shows the implantation scheme of patient 1. Signals were recorded using a standard clinical EEG system (XLTEK, subsidiary of Natus Medical) with 500 Hz of sampling rate, except for patient 3, where a sampling rate of 250 Hz was used (Fig. 1B). SEEG recordings from a total of 67 spontaneous seizures were collected and analyzed. Seizure onset and termination times of each seizure were independently marked by two epileptologists (A.P and R.R.). For each seizure we selected the marked ictal epoch together with 60 seconds of pre-ictal baseline activity. Artifacted channels were identified by visual inspection and removed prior to data analysis. A broadband-pass filter (FIR, filter band [1,165] Hz) was used to remove slow drifts and aliasing effects. We also used a notch FIR filter at 50 Hz and its harmonic frequencies to remove the power line interference.

#### Setting a ground truth: seizure onset zone and post-surgical outcome

After electrode implantation and monitoring, SOZ was identified by neurologists using visual inspection. In each patient, ictal activity was found to start in 3-14 channels that were marked as being part of the SOZ. In total, 78 recording sites were defined as the SOZ across patients, which accounts for 9% of the 906 implanted contacts. Surgical resection or radio-frequency thermocoagulation (RF-TC) was planned based on individual SEEG evaluations. At the time of writing (June 2019), Patients 1-3 had achieved seizure freedom after surgical resection (Engel I) with a follow-up period of 6 years. Patient 4 achieved seizure freedom after electrode explantation without need of resective surgery (Engel I) with a follow-up of 5 years. Patient 5 was initially responsive to RF-TC (Bourdillon et al., 2017), but after seizure relapse underwent resective surgery, after which achieved seizure freedom (Engel I) with a follow-up of 3 years. Patients 6, 7 and 8 underwent only RF-TC. Patients 6 and 7 are seizure free (Engel I) with a follow-up of 3 years. Patient 8 was initially responsive to RF-TC showing a seizure reduction larger than 50%, but relapse occurred 2 years after the procedure. Resective surgery was not an option for this patient due to cognitive risks. Patient 9 had achieved seizure freedom after surgical resection (Engel I) with a follow-up period of 5 years. We also included one patient (patient 10) in which the outcome was not successful (Engel III) because the brain resectomy failed to completely remove the identified seizure focus.

#### Statistical analysis

For each patient, the SOZ was defined by accumulating all seizure-specific detected SOZ regions. Then we used the SOZ defined by epileptologists as the ground truth to assess the performance of our method. For each patient, we computed the sensitivity and specificity of our selection. Additionally, we studied the effect of the thresholding procedure by computing the sensitivity and specificity of the method for a range of threshold values and extracted the area under the curve (AUC). This procedure was primarily done with SEEG signals in the monopolar configuration and with positive times. We then repeated the analysis using the bipolar configuration. We also assessed the amount of SOZ predictability carried in the pre-ictal activity, repeating the whole procedure with pre-ictal time windows (i.e., negative times from −30 to 0) in the monopolar setting. In all cases, we compared the area under the curve across settings to remove the threshold dependence of the results.

Finally, to relate the results of our method with validated post-operative information, we subselected patients that underwent resective surgery and had an Engel I post-surgical outcome (n=5). For this subset of patients, we defined three regions of interest: a) the seizure onset zone (SOZ), b) the resected zone (RZ), consisting of all recording sites that were surgically ablated, and c) the union of the two previous regions, consisting of all SOZ and/or RZ sites (SOZ+RZ) and therefore representing all putative sites with a critical role in the generation and/or spread of the patients’ seizure. For each of these regions of interest, we assessed the prediction power of the variable MA when considered only in seizure onset windows. In particular, we first normalized the MA values within each SOW to allow for cross-window comparison. For each patient, we averaged all normalized MA values across all SOWs of all seizures, thus defining a single normalized MA index (nMA) per patient’s site. By setting a cutoff value on the nMA index, we defined a binary classifier that was then compared to each region of interest. In each chase, the predictive value of the classifier was assessed by extracting the area under the curve (AUC) per patient.

### Data availability

The data that support the findings of this study are available from the corresponding author upon request.

## Results

### Magnitude and spread of enhanced oscillations in the peri-ictal period

In all studied patients and seizures, we computed all channels’ MA activation in the preselected frequencies and time windows of interest and extracted the GA and AE from the MA profiles. We first studied the extent to which both variables were providing redundant information. For each seizure, we computed the Pearson and Spearman correlation coefficients between the two variables. This computation yielded small-medium correlation values (Spearman’s r=-0.4±0.2, mean ± standard deviation across n=67 seizures), confirming that both variables were providing non-redundant information.

In this stage, we investigated the effect of the thresholding procedure used as part of the SOW detection algorithm. To this aim, we sought to characterize differences between the pre-ictal and ictal periods arising in the joint (AE,GA) empirical distribution at a group level (see the Supplementary Information for details). To do so, we normalized all GA values using the percentile score in each seizure and period (ictal, pre-ictal) and pooled all (AE,GA) values across seizures. A large number of ictal windows were found to cluster in the region with GA>80 and AE<0.5 (Supplementary Fig. 2A). On the other hand, pre-ictal windows exhibited a more sparse distribution of (AE,GA) values, with an average trend of higher AEs and lower GAs (Supplementary Fig. 2B). In particular, the region used for SOW detection in this study was GA>95 and AE<0.5 (See Materials and Methods). Then, for each seizure included in the study (a total of N=67) we compared the fraction of time-frequency windows satisfying both conditions with chance level (see the Supplementary Information for details). In both periods, the fraction of windows with GA>95 and AE<0.5 was found to be significantly above chance level (Supplementary Fig. 3), being this trend higher in the ictal period. Hence, we hypothesize that this association between very high and spatially confined activations is associated with ictal onset and constitutes an *a posteriori* validation of the rationale behind the procedure that we propose.

### Seizure-onset window detection

In 88% of the analyzed seizures (59 out of 67) the method was able to identify time-frequency windows satisfying the required criteria on the magnitudes of the variables GA and AE. In the remaining eight seizures no window fulfilled the conditions due to a number of reasons, including low spectral activations below the minimum required threshold of 3 standard deviations with respect to the pre-ictal baseline (1 seizure), widespread simultaneous activations that did not allow for a confined region to be safely identified (five seizures) or large pre-ictal activations comparable to the ictal activity itself (two seizures). The latter was the case of patient 5. Reviewing of the SEEG recordings from the two discarded seizures revealed that ictal onset was indeed preceded by continuous ictal-like activity.

### Seizure-onset windows unveil characteristic electrophysiological signatures

In each seizure, the SOWs were qualitatively found to pinpoint the characteristic frequency and time windows of the seizure onset patterns. As an example, in the first seizure of patient 1, the algorithm selected the following SOWs: 107-130 Hz during the first hundreds of milliseconds and 12-31 Hz between 10 and 20 s. Regions of the posterior and anterior hippocampi were selected as being inside the SOZ. Inspection of the electrophysiological activity recorded around seizure onset revealed that the output of the method was qualitatively describing the seizure onset patterns (Fig. 4A).

**Figure 4.**
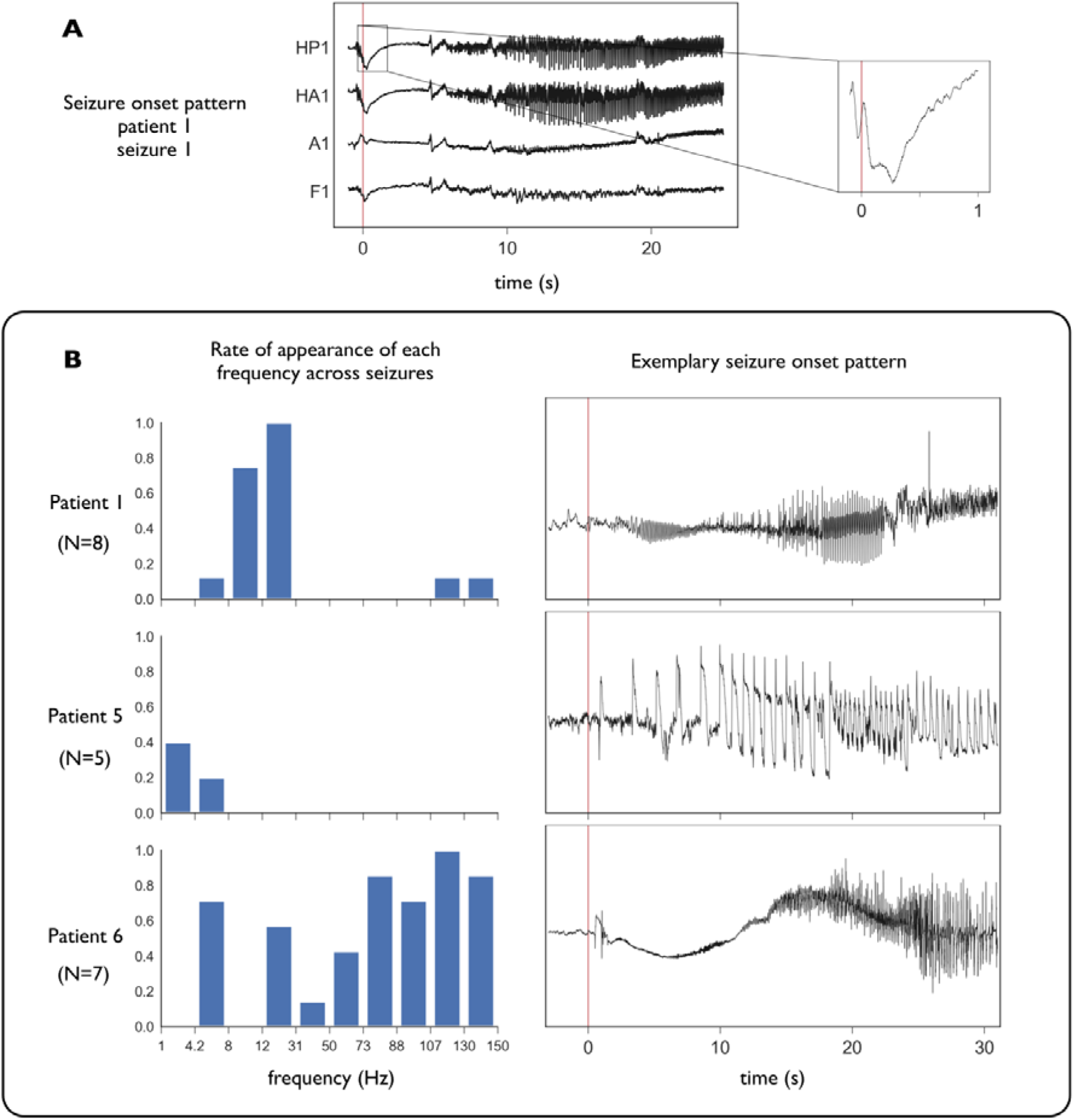
Seizure-onset windows unveil epileptogenic pattern signatures. (A) Detail of the seizure onset pattern in a seizure of patient 1. We show electrophysiological activity recorded at two SOZ contacts (HP1, HA1) and two contacts outside the SOZ (A1, F1). Seizure onset is marked with a red vertical line. As suggested in Fig. 3, the seizure is initiated at a hippocampal level with rapid discharges (∼110 Hz) of very low amplitude in the first hundreds of milliseconds that are particularly clear in HP1. This activity is followed by low-voltage fast-activity (LVFA) at 12-31 Hz that becomes visible 5 seconds after seizure onset and that increases in amplitude as the seizure progresses. Activations observed at 12-31 Hz are in fact a combined effect of LVFA activity (∼30 Hz) together with slow rhythmic spikes (RS) (∼15 Hz) of high amplitude particularly observable between 10 and 20 s. This activity then propagates into other regions and is no longer specific to SOZ regions (not shown). (B) Seizure onset frequencies along with their corresponding seizure onset patterns for three representative patients with N=8, 5 and 7 seizures, respectively. (Left) For each patient and frequency of interest, the bar height indicates the fraction of seizures in which the frequency appeared to be part of the SOW. (Right) In each case, we show the electrophysiological pattern recorded at an exemplary SOZ site after seizure onset for a given seizure. Seizure onset is marked with a red vertical line. In patient 1 seizures typically start with LVFA (∼30 Hz) combined with RS (∼10 Hz). In the example, rhythmic slow waves (RSWs) in the θ band (4-8 Hz) are also observed. In patient 5 seizure onset is characterized by very slow spiking activity (∼1 Hz) of large amplitude that becomes faster (up to ∼4 Hz) as the seizure progresses. Patient 6 has a much more spectrally diffuse activity (50-150 Hz) of very low amplitude. As seen in the example, there is also a very slow wave or baseline shift that was not detected by our method.

SOWs were found to be heterogeneous across patients and, in some cases, even across seizures of the same patient. For each time-frequency window of interest, we computed the fraction of seizures for which that window was a SOW (Fig. 4B). The most common SOWs were found to be in the frequency range 12-31 Hz with integration times spanning from 10 to 30 s after seizure onset. Around half of the seizures analyzed in this study had seizure onset patterns characterized by these time-frequency windows, roughly corresponding to low-voltage fast-activity (LVFA) combined with rhythmic spikes (RS) observed in the first seizure of patient 1. Other patterns at higher or lower frequencies and around 20 seconds after seizure onset were also observable. Of particular interest was the SOW at 107-130 Hz during the first milliseconds after seizure onset, which is consistent with the literature about HFOs being a good biomarker of pathological epileptic activity (Murphy *et al*., 2017).

Additionally, for each patient and frequency of interest, we computed the fraction of seizures in which that particular frequency appeared to be part of the SOW. Roughly, three different frequency distributions could be identified within our cohort. Fig. 4B shows these distributions for three exemplary cases and the seizure onset patterns they correspond to as described in (Lagarde *et al.*, 2016). In Patient 1 the most relevant frequencies were found to be between 4.2 and 12 Hz. In this case, seizure onset is characterized by LVFA + RS activity that becomes particularly visible around 10 s after seizure onset. In some cases, fast discharges (between 110 and 150 Hz) have also a role during the first milliseconds of the ictal period, while in others rhythmic slow waves (RSWs) in the θ band (4-8 Hz) can be observed. Seizure onset in patient 5 was found to be characterized by very slow spiking activity (∼1 Hz) of large amplitude in between 10 and 20 seconds after seizure onset, that becomes faster (up to ∼4-8 Hz) as the seizure progresses. Patient 6 exhibited a much more spectrally diffuse activity (ranging from 50 to 150 Hz in most seizures) of very low amplitude that becomes sufficiently high for SOZ discrimination around 20 s after seizure onset.

### Seizure focus prediction

For each patient, the SOZ was defined by accumulating all seizure-specific detected SOZ regions. Then we used the SOZ defined by epileptologists as the ground truth to assess the performance of our method. For each patient, we computed the sensitivity and specificity of our selection. In order to quantify the localization error, we also computed the average Euclidean distance between clinically marked SOZ contacts and the output of our analysis. Patient specific sensitivities and specificities are reported in Fig. 5. The average sensitivity of the method across patients was 0.94±0.03 (mean ± standard error of the mean), with an average specificity of 0.90±0.03 (mean ± standard error of the mean). In patients 1, 3, 5, 7, 8, 9 and 10 all SOZ regions were identified by our method (sensitivity=1). In the remaining cases (patients 2, 4 and 6) false negatives (SOZ contacts mistakenly marked as non-epileptogenic) lied at most 1 contact (i.e. 1.5 mm) apart from true positives (regions correctly marked as epileptogenic).

**Figure 5.**
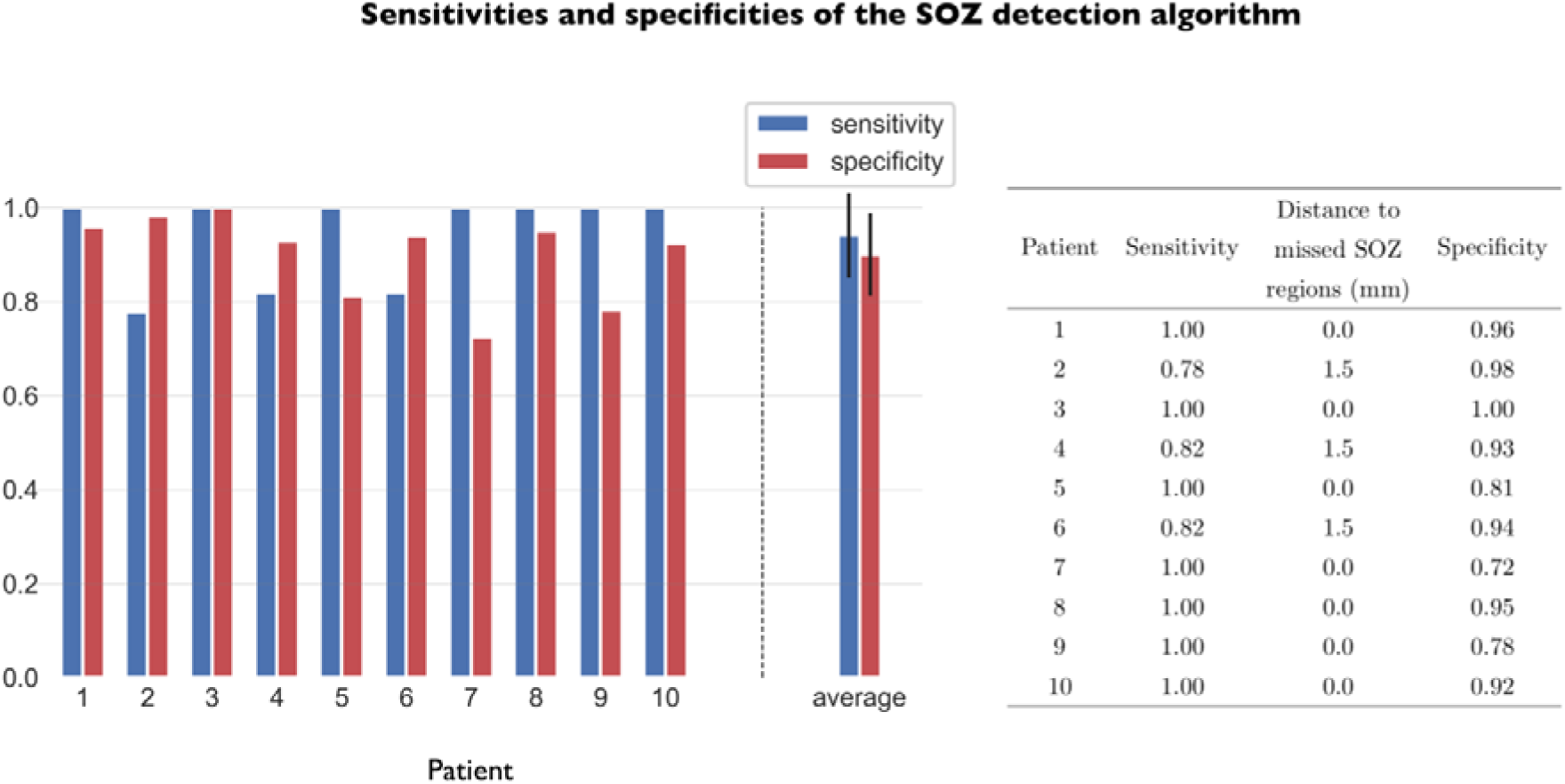
Validation of the method: SOZ detection accuracy. Automatically detected SOZ was compared with the clinically marked SOZ regions. The figure reports sensitivities and specificities at a subject and group levels. The average sensitivity of the method across patients was 0.94±0.03 (mean ± standard error of the mean). The average specificity was 0.90±0.03 (mean ± standard error of the mean). Only in patients 2, 4 and 6 the identification was not complete. However, false negatives (SOZ contacts mistakenly marked as non-epileptogenic) lied at most 1 contact (i.e. 1.5 mm) apart from true positives (regions correctly marked as being inside the SOZ).

### Stability analysis I: Comparative performance between monopolar and bipolar configurations

This procedure was primarily done with SEEG signals in the monopolar configuration and setting the AE and GA thresholds at 0.5 and 95-th percentile, respectively. Then, we investigated the influence of the signal referencing in the performance of the method. The whole analysis was repeated using the bipolar configuration. In order to compare the level of performance in the two settings independent of the selected algorithm working point we computed the patient-average sensitivity and specificity of the predicted SOZ as a function of the GA and AE thresholds in both configurations. Dependence on the two thresholds was tested independently of each other.

First, we computed the sensitivity and the specificity of the method as a function of the GA threshold (from 0 to 100) while keeping the AE threshold fixed around 0.5 (in the range 0.4-0.6). Then, we extracted the area under the curve (AUC) and compared it between SEEG configurations, representing the variability across the AE threshold with one standard deviation (see Fig. 6 for details). The same procedure was repeated to assess the performance as a function of the AE threshold (from 0.31 to 0.69) while keeping the GA threshold fixed around 95 (in the range 92-98).

**Figure 6.**
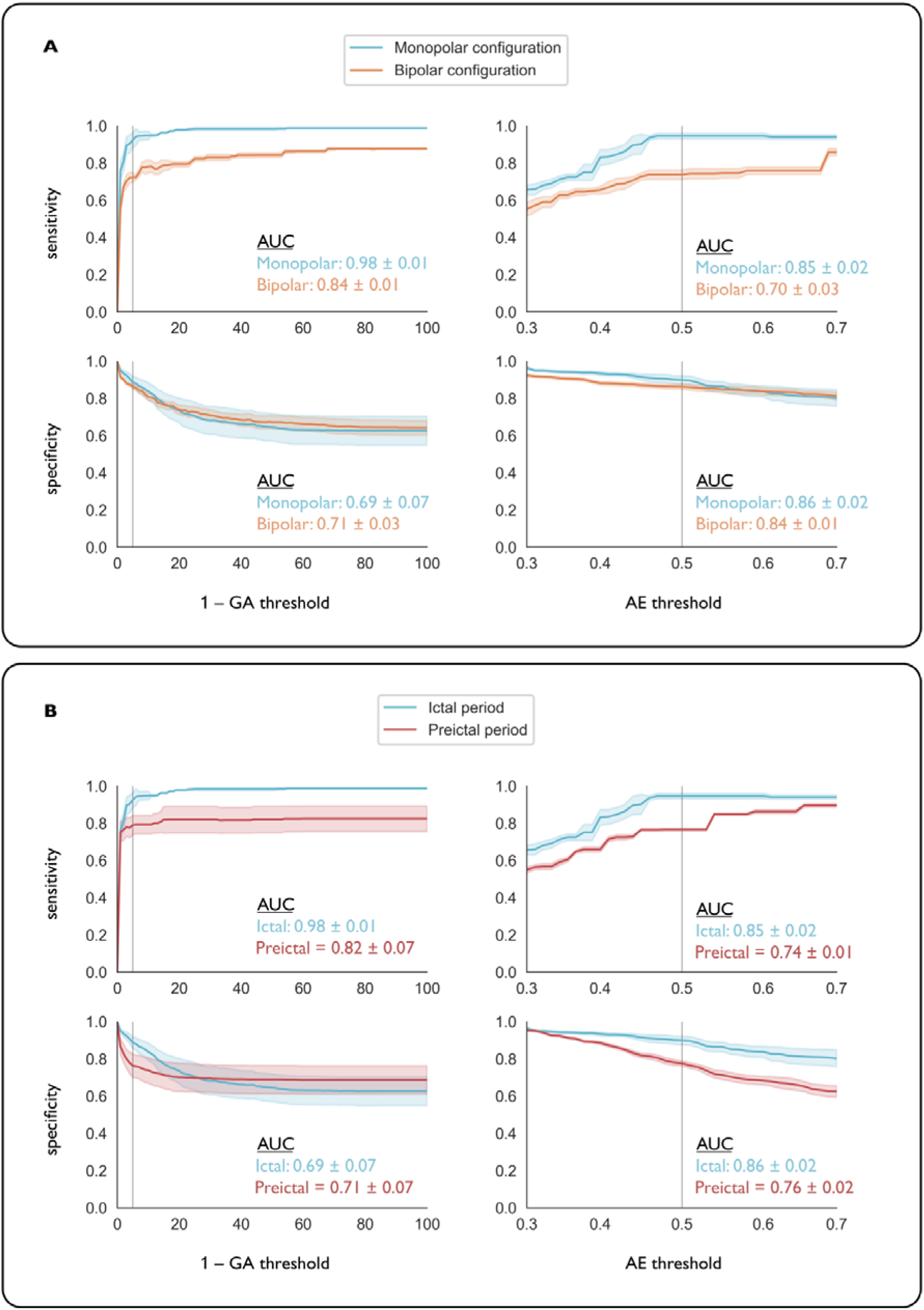
Patient-average performance is compared under different threshold conditions, recording configurations and time periods. Patient-average performance of the SOZ detection algorithm is shown as the thresholds are varied under different conditions: mono- vs bipolar referencing, and pre-ictal vs ictal periods. For each threshold type, variability is shown with respect to the other threshold type and represented by one standard deviation of the values. (Left graphs) For each SEEG configuration we show the sensitivity and the specificity of the method as a function of the GA threshold while keeping AE fixed around 0.5. In particular, for each value of the GA threshold, the solid line shows the average performance across all AE threshold values in the range 0.4-0.6 (in steps of 0.01, N=20), while the shaded area depicts one standard deviation across AE threshold values. We also report the area under the curve (AUC) relative to its theoretical maximum, i.e., divided by 100. In all cases, vertical lines mark the working point of the algorithm in our study (GA and AE thresholds set at 95 and 0.5, respectively). (Right graphs) For each SEEG configuration we show the sensitivity and the specificity of the method as a function of the AE threshold while keeping GA fixed around 95. The solid line shows the average performance across GA threshold values in the range 92-98 (in steps of 1, N=6), while the shaded area depicts one standard deviation across GA threshold values. AUC values are also reported relative to the theoretical maximum area of 0.4. (A) Performance comparison between mono- and bipolar referencing. While there is no significant difference in terms of specificity between monopolar and bipolar configurations, the sensitivity of the method is significantly higher in the monopolar configuration. (B) Performance comparison between pre-ictal and ictal time periods. Although the method has a better performance in the ictal period, high sensitivities and specificities indicate that the pre-ictal period contains sufficient information for SOZ prediction.

While no relevant differences were found in terms of specificity, the algorithm was found to be significantly more sensitive to SOZ regions in the monopolar configuration (Fig. 6A). In particular, the relative AUC of the sensitivity as a function of the GA threshold was found to be 0.98±0.01 in the monopolar configuration, while it dropped to 0.84±0.01 in the bipolar configuration. On the other hand, while varying the AE threshold the relative AUC of the sensitivity was 0.85±0.03 in the monopolar configuration and 0.70±0.03 in the bipolar setting.

### Stability analysis II: Focus prediction before seizure onset

Additionally, we assessed the amount of SOZ predictability carried in the pre-ictal activity. In order to do so, we repeated the same analysis described in the previous section using pre-ictal time windows (i.e., negative times from −30 to 0) in the monopolar setting. The results were compared with those obtained with positive times (Fig. 6B). The average sensitivities and specificities of the method across patients in the pre-ictal period were lower than in the ictal period: 0.77±0.32 and 0.77±0.12, respectively. Yet, although discrimination is lower in the pre-ictal period, these results prove that there is useful information to predict the SOZ already before seizure onset in most of the patients.

### Method validation with post-surgical outcome

For patients that underwent resective surgery and had a very good post-surgical outcome (Engel I, N=5), the binary classifier induced by nMA indices in the SOW selected by the algorithm was compared across the three regions of interest (SOZ, SOZ+RZ and RZ). ROC curves for each region of interest are shown in Fig. 7. SOZ prediction achieved very high AUC values: 0.997 ± 0.001 (mean ± standard error of the mean). When compared to RZ+SOZ and RZ, the classifier achieved lower but still notable predictive power values (0.79 ± 0.05 and 0.69 ± 0.08, respectively), indicating that the nMA variable carries relevant information not only for SOZ identification, but also for RZ prediction.

**Figure 7.**
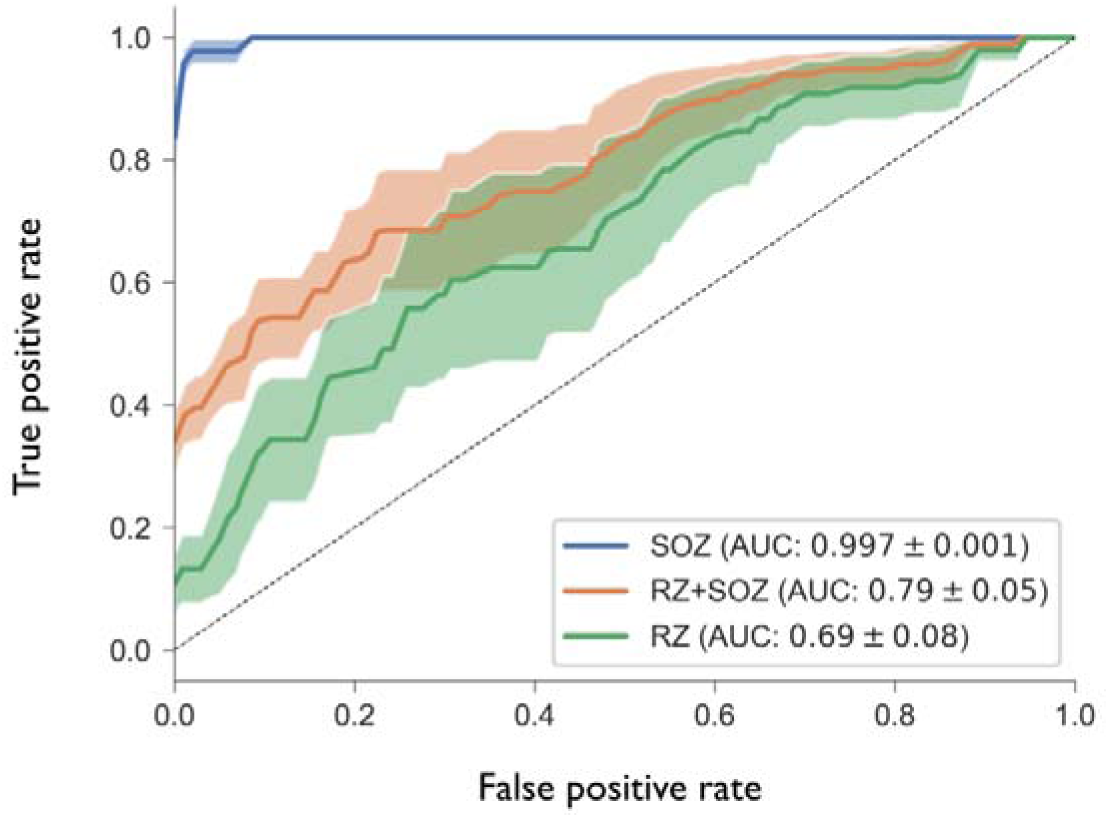
ROC curves for the prediction of SOZ and RZ regions in Engel I patients. For patients that underwent resective surgery and attained very good post-surgical outcome (Engel I, n=5), we computed the average nMA over all seizure onset windows, thus obtaining a single nMA index per site. We then computed the ROC curves when using these indices to predict the SOZ, RZ+SOZ and RZ regions, respectively. For each region, the solid line represents the average performance across patients, while the shaded area depicts the standard error of the mean. AUC values are also reported in the figure (mean ± standard error of the mean). The results suggest that the variable nMA carries relevant information for RZ prediction in patients that attained seizure freedom after resective surgery.

## Discussion

We proposed a novel methodology to automatically estimate the localization of spatially confined events in brain activity using the low entropy properties of emerging neural oscillations associated to those events. Our analysis was motivated by the problem of SOZ identification from intracranial EEG signals in patients with pharmacoresistant epilepsy. We tested the method performance with a cohort of ten patients with heterogeneous seizure-onset patterns and validated with the post-operative outcome of resective surgery.

Based on previous studies (Bartolomei *et al*., 2008; David *et al.*, 2011) we defined ictal-driven activity as an increase in signal power from pre-ictal to ictal epochs. In particular, we used the mean activation (MA) measure (Vila-Vidal *et al*., 2017), which quantifies the average spectral activation of each targeted brain structure for pre-defined frequency and time windows of interest (Fig. 1). However, while most studies constrained spectral activations to occur in preselected frequency bands, our method automatically infers the characteristic temporal scale and frequency range of locally enhanced oscillations in each case, thus maximizing the amount of information relevant for SOZ detection in a patient and seizure-specific context. Hence, by avoiding such frequency constraints, the method accounts for the intrinsic variability of initiation patterns (Lagarde *et al.*, 2016) and can flexibly adapt to the potential heterogeneities across different seizures of the same patient.

The proposed automated SOZ detection is performed using a two-stage procedure (Fig. 3). First, the most relevant frequency and time windows of interest (the so-called seizure onset windows, SOW) are extracted from the intracranial EEG signals. The algorithm relies on finding time-frequency windows in the transition between pre-ictal and ictal states that yield maximal and spatially confined spectral activations, thus ensuring that propagation has not started and that SOZ contacts can be naturally discriminated from other sites. Then, the most active channels are selected and accumulated over seizures to define the SOZ. The critical step of our analysis is the definition of two measures (Fig. 2), namely the global activation (GA) and the activation entropy (AE), that are later jointly optimized to find the relevant time-frequency window for SOZ localization. On one hand, GA is used to quantify maximal activations with respect to the pre-ictal basal state. Note that in the case where one contact has a very large MA compared to the others, the global activation coincides with the maximum of the MA distribution. However, the robustness of this measure makes it preferable when compared to a summary statistic that takes only one value. On the other hand, the AE is used to characterize how spread spectral activations are across recording sites, independent of the magnitude of these activations, a feature that is strictly measured by GA.

Although used with the mean activation (MA), the two measures and the optimization procedure that we described here are rather general and can be used in combination with other variables to find spatially confined events defined by changes in the magnitude of those variables. Specifically, the methodology that we developed may be useful in a number of settings to unravel the characteristic time and frequency scales (pattern signatures) of localized abrupt changes in brain activity during events of interests of both cognitive and clinical nature. Additionally, the method could also be used to extract the shape of the brain response to each of the considered stimuli.

We have shown that ictal onset is characterized by a general shift towards lower AEs and higher GAs. More specifically, we computed the density of (AE,GA) across seizures in the ictal period and pre-ictal period. A highly populated cluster defined by very high GAs and low AEs could be clearly identified in the ictal epoch. Based on this result, the method finds those windows with maximal global activations (large GA) under the constraint that these activations are spatially confined to a few regions (low AE). These windows are then used to reliably find SOZ sites (Fig. 3). Yet, the specific choice of the thresholds (GA above the 95-th percentile and AE<0.5) is rather arbitrary and deserves to be further discussed. We could have used a clustering algorithm to extract the specific boundaries of the dense cluster. However, this would yield more relaxed conditions that would result in a decrease in specificity. Hence, we chose to manually set the thresholds and tune them depending on the desired sensitivity and specificity of the output. The stability of the results around the chosen threshold values ensures the reliability of the method, making it robust to particular choices. Additionally, we did a surrogate analysis and compared the fraction of windows satisfying the required criterion with chance level. This analysis showed a strong correlation between the conditions both in the pre-ictal and ictal periods. We hypothesize that this correlation between very high and spatially confined activations is associated with ictal events and constitutes an *a posteriori* validation of the rationale behind the procedure that we propose. Moreover, a non-negligible consequence of the thresholding step is that seizures where SOZ discrimination cannot be guaranteed with a minimum level of confidence are discarded beforehand, thus ensuring meaningful SOZ detections. Discarded seizures can be further interpreted and might offer complementary insight during the pre-surgical evaluation. Although the treatment of such cases is out of the scope of this study, clinical reviewing of the SEEG recordings from patient 5 showed that the two seizures that had been rejected were indeed preceded by ictal-like activity.

A primary aspect of our analysis is that automatization does not come at the expense of interpretability, as it provides an output that can be fully read in clinically terms (Fig. 4). As seen in (Lagarde *et al.*, 2016) there is a wide range of seizure onset patterns and while some have a higher prevalence than others, high-frequency discharges cannot be considered to be the only epileptogenic onset biomarker. We have shown that the time-frequency windows selected by the method reliably characterize a variety of electrophysiological seizure onset patterns. In particular, we could isolate the high-amplitude low-frequency spiking activity at 1 Hz that defines the start of the ictal events in one of the analyzed patients.

SOZ detection was primarily done in the monopolar setting and using time windows from ictal epochs (Fig. 5). The average sensitivity across patients was 0.94±0.03, with an average specificity of 0.90±0.03 (mean ± standard error of the mean). SOZ detection was not complete only in three patients (2, 4 and 6) but missed SOZ sites lied at most 1.5 mm apart from the delineated region. We also studied the effect of the thresholding on the performance of the detection algorithm. The GA threshold (95th percentile) was found to lie at the onset of the sensitivity function stabilization, while in a range of high specificities. On the other hand, the AE threshold (0.5) lies on a plateau of the performance function. As the conditions on GA and AE are relaxed more non-SOZ sites are chosen. Despite being outside the SOZ, we hypothesize that these sites might have a critical role in sustaining and propagating epileptic activity in the early stages of the seizure.

Complementary to the main results, we further investigated the effect of the recording referencing on the method performance (Fig. 6A). While no significant differences were found in terms of specificity, the algorithm was more sensitive to SOZ regions in the monopolar configuration. Very few works have assessed the effect of the recording reference on intracranial EEG analysis and they have mainly studied its impact on connectivity measures (Arnulfo et al., 2015). To the best of our knowledge, this is the first study to compare the performance between monopolar and bipolar configurations for SOZ detection. The monopolar referencing yields data contaminated by volume conduction and remote field effects, while the bipolar montage averages out these effects offering a more localized spatial resolution. Yet, we hypothesize that in the case of a partially mapped SOZ, the monopolar configuration might be more sensitive to capture the activity from the missed SOZ sites and thus perform better when used to approximate the focus localization.

Additionally, we chose to explore the performance of the method in a short pre-ictal period (Fig. 6B). Although the method has proven to have a higher performance in the ictal period, the results bring evidence that the pre-ictal period also carries information that might be of interest for SOZ localization, as already seen in previous studies (Andrzejak et al., 2015; Tauste Campo et al., 2018).

Post-operative validation in Engel I patients revealed the potential predictive power of the variable nMA as a biomarker of the resected zone when averaged across seizure onset windows (Fig. 7). This result suggests that the features upon which the method is built characterize the generation, spread and maintenance of epileptic seizures. Yet, an average predictive value (AUC) for the RZ of 0.69 ± 0.08 (mean ± standard error of the mean) highlights the non-trivial relationship between the SOZ and the whole epileptogenic zone, in line with previous studies (Lüders et al., 2006; Huang et al., 2012; Rummel et al., 2015; Geier et al., 2015).

To sum up, the method that we proposed achieved very high accuracy values when used to predict SOZ regions from peri-ictal SEEG recordings. Our approach relies on the detection of the seizure onset time defined by the clinical neurophysiologists. Hence, in combination with automated seizure onset algorithms, it could be easily integrated as a complementary diagnostic tool with minimal computational costs for surgical planning, reducing time-consuming SEEG revisions and improving the clinician decision after pre-surgical evaluation.

The number of patients satisfying sufficiently long post-operative follow-up periods (10) constitutes the major limitation of this study. Further research should be conducted to validate our algorithm with a larger cohort of patients and beyond temporal lobe epilepsy. Additionally, the proposed procedure should be tested in a variety of alternative settings that involve the study of alterations in brain activity that are consistently time-locked to an event (Luck et al., 2000; Lauchaux et al., 2012; Kropotov, 2016). Overall, our methodology provides a robust computational approach that can be used to identify neural populations undergoing emerging functional or pathological transitions in settings of both experimental and clinical relevance.

## Acknowledgments

M.V. is supported by “la Caixa” Foundation 100010434 (LCF/BQ/DE17/11600022). G.D. is supported by the Spanish Ministry Research Project PSI2016-75688-P (AEI/FEDER), by the EU H2020 FET Flagship Human Brain Project 785907 (HBP SGA2), by the Catalan Research Group Support 2017 SGR 1545, and by the Swiss National Science Foundation (CRSII5_170873). We thank Mar Moya Giménez for her contribution to the initial stages of this work.

