## Supplementary Information for "Low entropy map of brain oscillatory activity identifies spatially localized events: a new method for automated epilepsy focus prediction"

#### 1 Methods

##### 1.1 SEEG signal analysis

The methodology developed in this study relies on finding the frequencies and time windows of interest where ictal activity is maximal with respect to a baseline pre-ictal period, while being spatially confined to a few contacts. Central to this approach is the definition of two novel measures, namely the global activation (GA) and the activation entropy (AE), that are used to monitor the magnitude of spectral changes with respect to the pre-ictal epoch and the spread of these spectral activations, respectively, at different frequencies and as time progresses from seizure onset. By setting appropriate conditions on the two measures it is possible to find time-frequency windows of interest where SOZ regions can be optimally discriminated. These windows will be closely linked to the spectral properties of each patient's electrophysiological seizure onset patterns and will therefore be called seizure onset windows (SOW).

Our method makes use of the mean activation (MA), a measure that was proposed in a recent study (Vila-Vidal et al., 2017) to quantify average spectral activations of targeted brain structures in a given frequency and time window of interest. In this section, we first review the methodological procedure that leads to the computation of each region's MA for a given time-frequency window. Then, we define the two core measures of the study, the global activation and the activation entropy, and discuss some of their fundamental properties. Based on these measures, in the last section we show how to find the seizure onset windows (SOW) and how to delineate the seizure onset zone (SOZ).

###### Mean activation (MA) in a time-frequency window of interest

The mean activation (MA) measures the average spectral activation of each targeted brain structure for pre-defined frequency and time windows of interest (Vila-Vidal et al., 2017). MAs are estimated from SEEG signals with a two-step procedure. Fig. 1C illustrates how the signal is processed to obtain the MAs in a given time-frequency window of interest. First, time-dependent spectral activations in the frequency of interest are found using the Hilbert transform method. All contacts' signals are band-pass filtered in non-overlapping logarithmically spaced narrow frequency bands $[f, f + \Delta f]$ (we used $\Delta f = 0.1$) covering the whole frequency of interest. Signal power in each narrow band is obtained by squaring the signal envelope (modulus of the analytical signal) and summation over all narrow bands is performed to obtain the time-dependent power of each region's SEEG signal in the desired frequency of interest. The resulting values are z-scored with respect to a baseline distribution defined by the power values of all contacts during the first 40 seconds of the pre-ictal period to obtain normalized power time courses per contact that can be interpreted as a measure of the power change during the ictal event.

For the precise estimation of time-average spectral activations, artifact-induced noise must be first removed. Time-stamps with frequency-specific artifacts simultaneously affecting the majority of SEEG signals are detected with a sliding-window analysis (200 samples width, 1 sample step), performed independently in the pre-ictal and ictal epochs. Time windows where the product of the mean correlation and the contact-average signal power is two standard deviations larger than the median are considered as artifacts and are discarded in the subsequent analysis. Finally, all power values are averaged in the time window of interest to obtain the mean activation MA of each monitored brain region for the selected time-frequency window of interest.

##### Assessing peri-ictal information with Global activation (GA) and Activation entropy (AE)

The criterion used to find time-frequency windows that yield optimal SOZ detection derives from the fundamental observation that these windows should fulfill two conditions: a) that the SOZ contacts have become sufficiently active in the selected frequency band and time window and b) that the abnormal epileptic activity has not yet propagated to other brain regions. In brief, to ensure optimal focus detection we must ensure that there is a hierarchical and selective activation of SOZ contacts only. To this aim we define two novel measures that allow to monitor these conditions from the structure of the MA distributions. Fig. 2 illustrates how these measures are computed and used to assess the amount of information carried in each window of interest. In particular, a) the global activation (GA) quantifies the overall activation with respect to the pre-ictal basal state and b) the activation entropy (AE) quantifies how spread spectral activations are across recording sites. The question of finding the SOWs can then be translated into an optimization problem in which GA must be maximized under the constraint of a low AE.

In what follows we offer detailed definitions of these measures and discuss their fundamental properties. For the sake of clarity, we will denote the MA of a given region j in the frequency band f and computed over a time window spanning from the seizure onset until time t with the following notation: $MA_j(f,t)$. On one hand, the GA measures the magnitude of the most relevant spectral activations from the MA profile for a given time-frequency window of interest. It is defined as the weighted average MA over all contacts, where each contact's contribution is weighted by its own activation, thus ensuring that most active regions have a higher impact on the final value (Fig. 2A):

$$\mathrm{GA}(f,t) = \frac{\sum_{j=1}^{N} w_j * \mathrm{MA}_j(f,t)^2}{\sum_{j=1}^{N} w_j(f,t)},$$

with $w_j = \mathrm{MA}_j(f,t)$ if $\mathrm{MA}_j(f,t) > 0$, and $w_j = 0$, if $\mathrm{MA}_j(f,t) \leq 0$. On the other hand, the AE is defined as the entropy of the MA distribution and characterizes the spatial spread of spectral activations for the given time-frequency of interest. First, the MA histogram is computed using 10 bins homogeneously spaced between the minimum and maximum MA values. Probability values for each bin ($p_i$ for $i = 1, \ldots, 10$) are found as the fraction of contacts lying within the corresponding MA bin (Fig. 2A). We then compute the Shannon's entropy of this distribution using the formula:

$$\mathrm{AE}(f,t) = -\sum_{i=1}^{10} p_i \log(p_i).$$

As the binning is defined to cover the entire MA distribution, only strictly positive values are possible. AE ranges from values close to 0 (corresponding to spatially confined spectral activations) to a maximum value of log(10) $\approx$ 2.3 (corresponding to uniform activations).

In order to systematically explore different windows for optimal SOZ detection we first divided the frequency spectrum into 10 non-overlapping bands using the following cutting points: 1, 4.2, 8, 12, 31, 50, 73, 88, 107, 130 and 150 Hz. In each band we considered a set of 75 different integration time windows obtained by fixing the initial bound at seizure onset and varying the final bound from 0.1 to 30 s (in steps of 0.1 s in the range 0.1-5 s and steps of 1 s in the range 5-30 s). Additionally, we also considered time windows spanning from seizure onset into the past, i.e., with final bound at seizure onset and initial bound ranging from 30 s to 0.1 s before seizure onset, with the same spacing as before. For each time-frequency window, we computed the MA profile and extracted its GA and AE. Fig. 2B and 2C show the GA and AE distributions for one exemplary seizure of patient 1, respectively.

##### Maximizing peri-ictal information for automatic seizure onset zone (SOZ) localization

The procedure described in the previous section was sequentially applied to the 67 seizures included in this study. Fig. 3 summarizes the processing steps. For each patient and seizure, SEEG signals in the peri-ictal period were band-pass filtered in pre-defined bands of interest spanning the whole spectrum (Fig. 3A). Then, for each band, MAs were obtained for all possible of time windows of interest (Fig. 3B). For each time-frequency window, we extracted the GA and AE from the MA distribution (Fig. 3C). SOW detection was achieved by finding time-frequency windows that maximized the GA under the constraint of low AE to ensure that spectral activations were confined only to a few contacts. To find the SOWs we considered all pairs $(f, t)$ with positive integration times (i.e., excluding time windows in the pre-ictal state) and set two threshold conditions, one per variable. For each seizure, we first pooled together all GA values and set a GA threshold at the 95-th percentile of the distribution. In any case, the threshold was always kept larger than 3 to ensure significant global activations with respect to the pre-ictal state. On the other hand, as the AE measures the diversity of spectral activations, setting a threshold on this measure is equivalent to requiring a minimum fraction of regions to lie in the same MA histogram bin (see section 2 of the Supplementary information). In particular, the lower the entropy threshold, the higher the number of regions with similar MAs. The curve shown in Supplementary Fig. 1 gives a lower estimate of the fraction of brain regions with similar MAs for each AE value and can therefore be used to tune the entropy threshold. In this study, we used a threshold of 0.5, which requires at least 80% of the contacts to lie within the same MA bin (specifically, they were required to lie within the lowest MA bin).

All time-frequency windows satisfying both conditions were preselected as candidates to be SOWs. Finally, for each frequency band we kept only the first time windows satisfying the condition. Once the condition was violated, we assumed that propagation had occurred and discarded all subsequent time windows, regardless of their GAs and AEs. Fig. 3D shows the selected SOWs for the first seizure of patient 1. Since the SOWs were defined to guarantee spatially confined spectral activations, the identification of the SOZ followed naturally from the MA distribution. By definition, for each pair SOW, at least 80% of the contacts are confined to the first bin of the MA histogram, exhibiting very low activations. The few remaining active contacts were considered to be part of the SOZ (Fig. 3E). This procedure was repeated for all selected SOWs, and SOZ contacts were accumulated, thus obtaining a single SOZ per seizure (Fig. 3F).

#### 1.2 Setting a threshold on the Activation entropy

The activation entropy (AE) measures the diversity of spectral activations (MA distribution) across brain sites. Therefore setting an upper threshold on AE implicitly sets a condition on the spread of the MA distribution. Intuitively, the lower the entropy threshold, the higher the number of regions with similar MAs.

In particular, given an upper bound on the entropy of AE, lower bounds on probabilities (i.e., fractions of channels in the same bin) can be estimated using the entropy curve of a Bernoulli distribution. A Bernoulli distribution $\mathcal{B}(p)$ is a discrete random variable with two possible outcomes with probabilities $p$ and $1-p$, respectively. The entropy of a Bernoulli distribution is given by the following formula: $h_2(p) = -p\log p - (1-p)\log(1-p)$, for $p \in [0,1]$. The function $h_2$ is symmetric around $\frac{1}{2}$ and invertible when restricted to $p \in \left[\frac{1}{2}, 1\right]$ (Supplementary Fig. S1). Its inverse function $h_2^{-1}$ gives a lower estimate of the fraction of brain regions with similar MAs for each AE value and can be used to tune the entropy threshold.

The fundamental property that we use to extract a lower bound on the probabilities of particular AE bins can be formulated as follows. If the AE is smaller than a certain value $h$, which in turn is the entropy of a Bernoulli distribution $\mathcal{B}(p)$, for a certain value of $p$, then one of the 10 bins in the MA histogram must have a probability larger than $p$. This section aims to give the reader a general formulation and an explicit proof of this property of the entropy function.

##### General formulation

Let $X$ be a discrete random variable with possible outcomes $\{x_1, \ldots, x_n\}$ and probability mass function $P$. Let $p_i = P(X = x_i)$. The Shannon's entropy of $X$ is defined as

$$H(X) = -\sum_{i=1}^{n} p_i \log p_i. \tag{1}$$

In the case of $p_i = 0$ for some $i \in \{1, \ldots, n\}$, the value of the corresponding summand $0\log 0$ is taken to be 0, which is consistent with the limit:

$$\lim_{p \to 0+} p\log(p) = 0. \tag{2}$$

If we define the function $\eta(p) = -p\log p$, for $0 < p \leq 1$, and $\eta(0) = 0$, the entropy of $X$ can be rewritten as

$$H(X) = \sum_{i=1}^{n} \eta(p_i). \tag{3}$$

The function $\eta$ is continuous in $[0,1]$.

**Proposition 1.** *If $p > q$, and $0 \leq \Delta \leq q$, then:*

$$\eta(p) + \eta(q) > \eta(p+\Delta) + \eta(q-\Delta).$$

*Proof.* The function $f(\Delta) = \eta(p+\Delta) + \eta(q-\Delta)$ is continuous in $[0,q]$ and its derivative with respect to $\Delta$ has the following expression:

$$f'(\Delta) = \eta'(p+\Delta) - \eta'(q-\Delta) = -\log(p+\Delta) - 1 + \log(q-\Delta) + 1 = \log\left(\frac{q-\Delta}{p+\Delta}\right). \tag{4}$$

If $p > q$, then $p+\Delta > q-\Delta$ and $f'(\Delta) < 0$ for all $0 < \Delta < q$. Hence, the function $f$ is monotonically decreasing in $(0,q)$, which proves the proposition.

□

In particular, for $\Delta = q$, we have $\eta(p) + \eta(q) > \eta(p+q)$.

**Proposition 2.** *Let $h_2(p) = -p\log p - (1-p)\log(1-p)$ be the entropy of a Bernoulli distribution with parameter $p \in A = \left[\frac{1}{2}, 1\right]$. If $H(X) \leq t \in h_2(A)$, then $p_i \geq h_2^{-1}(t)$, for some $i \in \{1, \ldots, n\}$.*

*Proof.* Let us assume that all probabilities are smaller than $\tau = h_2^{-1}(t)$ and that they are indexed in increasing order ($p_0 < p_1 \leq \cdots \leq p_n < \tau$, with $p_0 = 0$). Let $j \in \{1, \ldots, n-1\}$ be such that

$$p_n + p_0 + \cdots + p_{j-1} \leq \tau < p_n + p_0 + \cdots + p_{j-1} + p_j. \tag{5}$$

By sequentially applying the property that $\eta(p) + \eta(q) > \eta(p+q)$ we obtain that

$$\begin{aligned} H(X) &= \eta(p_n) + \eta(p_1) + \cdots + \eta(p_{j-1}) + \eta(p_j) + \eta(p_{j+1}) + \cdots + \eta(p_{n-1}) > \\ &> \eta(p_n + p_1 \cdots + p_{j-1}) + \eta(p_j) + \eta(p_{j+1} + \cdots + p_{n-1}). \end{aligned} \tag{6}$$

Let $\Delta = \tau - (p_n + p_0 + \cdots + p_{j-1})$. From equation 5, $0 \leq \Delta < p_j$. Using P1:

$$\begin{aligned} H(X) &> \eta(p_n + p_1 \cdots + p_{j-1} + \Delta) + \eta(p_j - \Delta) + \eta(p_{j+1} + \cdots + p_{n-1}) > \\ &> \eta(p_n + p_1 \cdots + p_{j-1} + \Delta) + \eta(p_j - \Delta + p_{j+1} + \cdots + p_{n-1}) = \\ &= \eta(\tau) + \eta(1-\tau) = h_2(\tau) = t, \end{aligned} \tag{7}$$

which contradicts $H(X) \leq t$. Hence, the assumption that all probabilities are smaller than $\tau$ must be rejected.

□

### 2 Results

#### 2.1 Extracting peri-ictal information with the global activation and the activation entropy

In all studied patients and seizures, we computed all channels' MA activation in the preselected frequencies and time windows of interest and extracted the GA and AE from the MA profiles. We first studied the extent to which both variables were providing redundant information. For each seizure, we computed the Pearson and Spearman correlation coefficients between the two variables. This computation yielded small-medium correlation values (Spearman's $r = -0.4 \pm 0.2$, mean $\pm$ standard deviation across n=67 seizures), confirming that both variables were providing complementary information.

In this stage, we investigated the effect of the thresholding procedure used as part of the SOW detection algorithm. To this aim, we sought to characterize differences between the pre-ictal and ictal periods arising in the joint (AE,GA) empirical distribution at a group level. Prior to the computation of the density functions, we normalized all GA values using the percentile score in each seizure and period (ictal, pre-ictal) separately. We then pooled all (AE,GA) values across seizures and used kernel-density estimation with Gaussian kernels and automatic bandwidth determination as given by Scotts rule (Scott, 2015). Supplementary Fig. 2 shows the density of windows in each period. After seizure onset (Supplementary Fig. 2A), a large number of windows was found to cluster in the region with GA$>$ 80 and AE$<$ 0.5, a subset of which is targeted by our method. On the other hand, the 30 s preceding seizure initiation (Supplementary Fig. 2B) were characterized by a more sparse distribution of (AE,GA), with higher AEs and lower GAs on average.

More specifically, the region which is primarily used for SOW detection is defined by GA$>$ 95 and AE$<$ 0.5. For each seizure included in the study (a total of N=67) we compared the fraction of time-frequency windows satisfying both conditions with chance level, which was computed assuming no correlation between GA and AE. In this case, the fraction of time-frequency windows satisfying both conditions would be the product of the fraction of seizures satisfying each condition separately. We used a t-test to examine statistical differences between the two distributions and the fraction of windows within that subset was found to be significantly above chance level both in the ictal (N=67, $P < 10^{-9}$) and pre-ictal (N=67, $P < 10^{-7}$) periods (Supplementary Fig. 3), suggesting a strong correlation between the conditions set upon the two variables. We hypothesize that this correlation between very high and spatially confined activations is associated with ictal onset and constitutes an a posteriori validation of the rationale behind the procedure that we propose.

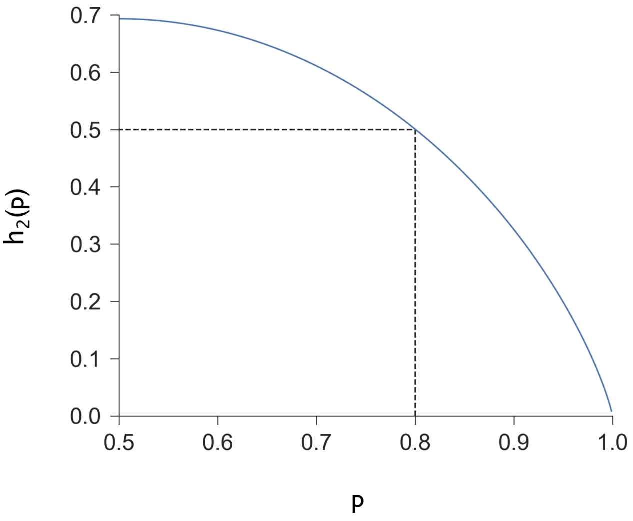


Figure S1: Entropy of a Bernoulli distribution $X \sim \mathcal{B}(p)$, with $p \geq 0.5$. The figure shows the entropy of $X$ as a function of the parameter $p$. Analytically, $\mathcal{H}(X) = \Phi(p) = -p \log p - (1-p)\log(1-p)$.

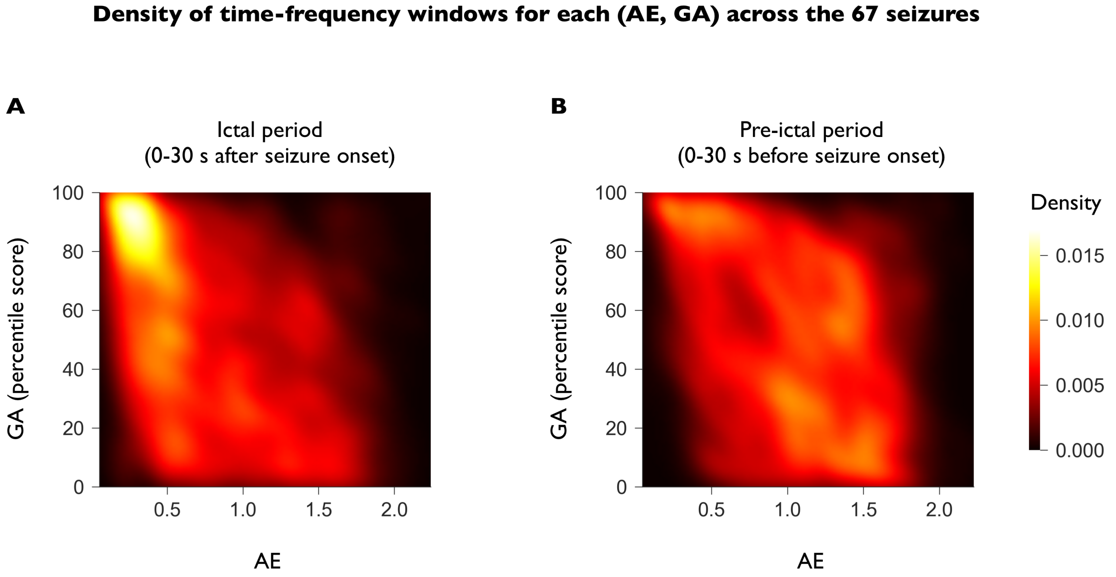


Figure S2: Distribution of AE and GA across the 67 seizures included in the study. (A) Density of time-frequency windows with given (AE,GA) across seizures in the ictal period (0-30 s after seizure onset). To compute the probability density function of the variable (AE,GA) we used kernel-density estimation with Gaussian kernels and automatic bandwidth determination as given by Scotts rule [Scott, 2015]. The plot shows an organized structure with a highly populated cluster in the region with GA$>80$ and AE$<0.5$, a subset of which if targeted by our method. (B) Density of time-frequency windows with given (AE,GA) across seizures in the pre-ictal period (0-30 s before seizure onset). The pre-ictal period exhibits a more sparse distribution of (AE,GA) values than the ictal period, with higher AEs and lower GAs on average.

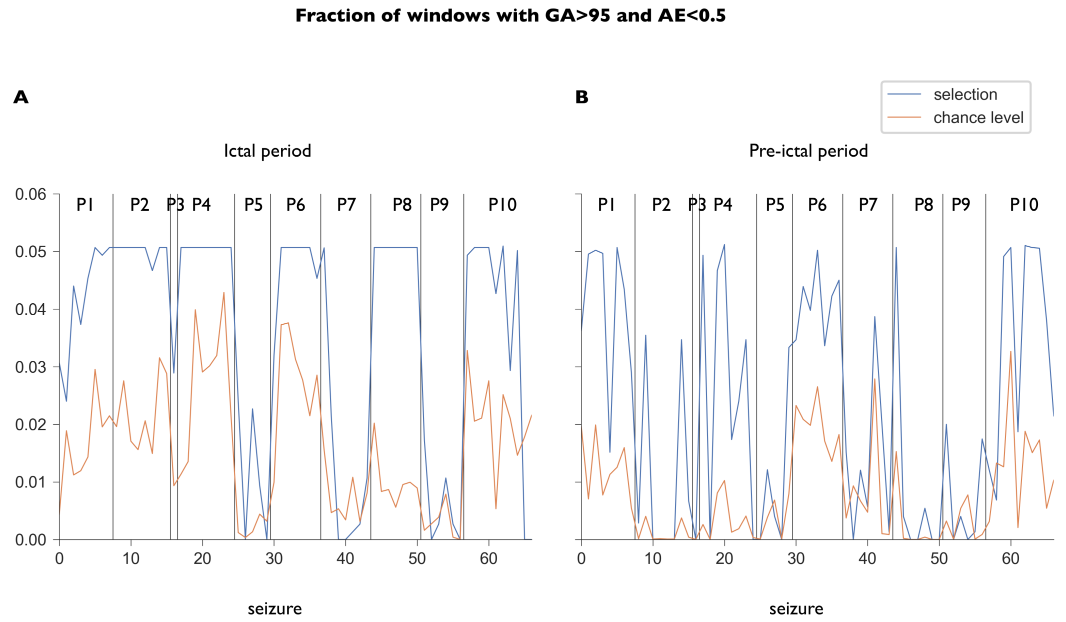


Figure S3: Relationship between the two thresholds. For each seizure (a total of N=67), the blue curves show the fraction of time-frequency windows with GA above the 95-th percentile and AE below 0.5 in the ictal (A) and pre-ictal (B) periods. Chance level was computed analytically for each period and seizxure seizure assuming no correlation between GA and AE. In this case, the fraction of time-frequency windows satisfying both conditions would be the product of the fraction of seizures satisfying each condition separately. Differences between the two distributions were examined using a t-test for independent samples and were found to be statistically significant both in the ictal ($N = 67$, $P < 10^{-9}$) and pre-ictal ($N = 67$, $P < 10^{-7}$) periods, suggesting a correlation between the two conditions. We hypothesize that this correlation between very high and spatially confined activations is associated with ictal onset and constitutes an a posteriori validation of the rationale behind the procedure that we propose.

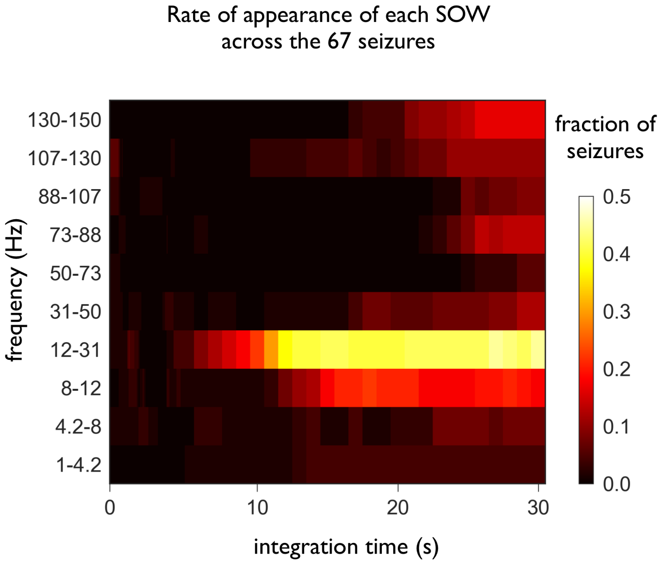


Figure S4: Rate of appearance of each SOW across the 67 seizures included in the study. The most common SOWs are in the frequency range 12-31 Hz with integration times spanning from 10 to 30 s after seizure onset. Around half of the seizures analyzed in this study have seizure onset patterns characterized by these time-frequency windows, roughly corresponding to the LVFA + RS activity already described in the first seizure of patient 1. Note, however, that other patterns at higher or lower frequencies and around 20 seconds after seizure onset are also observable. Of particular interest is the SOW at 107-130 Hz during the first milliseconds after seizure onset, which is consistent with the literature about HFOs being a good biomarker of pathological epileptic activity.
